# Spatial, temporal and molecular dynamics of swine influenza virus-specific CD8 tissue resident memory T cells

**DOI:** 10.1101/2021.08.23.457377

**Authors:** Veronica Martini, Matthew Edmans, Simon Gubbins, Siddharth Jayaraman, Basu Paudyal, Sophie Morgan, Adam McNee, Théo Morin, Pramila Rijal, Wilhelm Gerner, Andrew K. Sewell, Ryo Inoue, Mick Bailey, Timothy Connelley, Bryan Charleston, Alain Townsend, Peter Beverley, Elma Tchilian

**Affiliations:** The Pirbright Institute, Pirbright, UK; The Roslin Institute, Edinburgh, UK; Division of Infection and Immunity, Cardiff University School of Medicine, Cardiff, UK; Weatherall Institute of Molecular Medicine, University of Oxford, Oxford; Laboratory of Animal Science, Setsunan University, Osaka, Japan; Bristol Veterinary School, University of Bristol, Langford, UK; National Heart and Lung Institute, St Mary’s Campus, Imperial College, London, UK

## Abstract

We defined naïve, central memory, effector memory and terminally differentiated porcine CD8 T cells and analyzed their phenotype in lymphoid and respiratory tissues after influenza infection or immunization using peptide-MHC tetramers of three influenza nucleoprotein (NP) epitopes. The hierarchy of response to the three epitopes changes during the response in different tissues. Most NP-specific CD8 T cells in broncho-alveolar lavage (BAL) and lung are tissue resident memory cells (TRM), that express CD69 and have an effector memory or terminally differentiated phenotype. NP-specific cells isolated from BAL express genes characteristic of TRM, but gene expression differs at 7, 21 and 63 days post infection. The frequency of NP-specific cells declines over 63 days in all tissues but is best maintained in BAL. The pig is a powerful model for understanding how best to induce and harness local immunity to respiratory viruses.

**One sentence summary:** Influenza NP-specific porcine tissue resident memory CD8 T cells persist in the lung with major changes in gene expression.

## Introduction

Immunity to influenza A viruses (IAV) has been intensively studied over many years and the role of neutralizing antibody in protection against homologous virus is well established (reviewed (*1*)). Moreover experimental studies in mice and humans have also revealed the importance of pre-existing cellular immunity in heterosubtypic protection (reviewed (*2*)). The development of live attenuated influenza virus vaccines administered by nasal spray, now commercially available for humans and pigs, attests to the realisation that many lymphocytes reside in non-lymphoid tissues and that local immune responses are important in protective immunity (*3, 4*).

Intravenous infusion of anti-lymphocyte antibodies prior to isolation of lymphocytes from tissues has led to the definition of populations of lymphocytes that are considered to be tissue resident because they are not stained by the infused antibody. The majority of these cells have an activated or memory phenotype, indicating that they have most likely encountered antigen and they are therefore termed tissue resident memory cells (TRM) (*5–10*). In mice TRM have been shown to exceed the number of T cells in the lymphoid system and to play important roles in maintaining local immune memory (*11*). TRM are the predominant population in the adult human lung and antigen-specific cells were found at stable frequencies years after pathogen encounter, indicating their key role in respiratory infections (*12*). Mouse and human TRM express CD69 and more variably CD103, both essential for tissue retention (*6, 13–16*).

Porcine physiology closely resembles that of humans, pigs have a similar distribution of sialic acid in their respiratory tract and are infected with similar influenza viruses, making them a powerful natural host, large animal model to study immunity to influenza (*17–19*). Furthermore, they are an animal reservoir that poses a zoonotic threat to humans and influenza viruses are also a cause of significant economic losses to farmers (*20*). We have demonstrated the similarity of porcine and human antibody responses to influenza viruses, emphasized the importance of respiratory tract local immune responses in protective immunity and indicated that there are high frequencies of influenza-specific T cells in broncho-alveolar lavage (BAL), lung tissue and tracheo-bronchial lymph nodes (TBLN) (*21–24*). However, the lack of monoclonal antibodies (mAbs) to porcine CD69 and CD103 has made studies of porcine TRM challenging although we have previously identified porcine TRM, following infusion of anti-CD3 antibody (*23, 25*). Nevertheless, their phenotype and function during influenza infection remain poorly defined and there have been few *ex vivo* studies of antigen-specific T cells. In addition, while the phenotype of porcine helper T cells has been thoroughly analyzed (*26–29*), CD8 T cells are less well characterized and CCR7, essential for migration to lymphoid organs and CD45RA, a known differentiation marker, both widely used in human immunology, have never been studied in combination to define subsets of porcine T cells.

Here we examined influenza virus-specific CD8 TRM throughout the respiratory tract but focussed on CD8 TRM in BAL as this population of airway cells is relatively accessible in many species including humans, contains almost exclusively TRM apart from alveolar macrophages and does not require extraction procedures that might alter the cellular composition and phenotype. Here, for the first time we showed that CD45RA and CCR7 together identify porcine CD8 T cell subsets similar to those in humans and we described the expression of CD69 in CD8 T cells in several tissues, using a newly generated antibody (*30*). We examined antigen-specific CD8 TRM in the context of influenza infection and immunization in inbred Babraham pigs (*31*), using peptide-SLA tetramers carrying a previously identified (*32*) and two novel influenza nucleoprotein (NP) epitopes. This enabled us to define the phenotype of influenza-specific CD8 T cells, analyze their distribution in different tissues, define their transcriptional profile at different times after infection and model the dynamics of the response.

## Results

### Subsets of porcine T cells and identification of CD8 TRM

Antibodies to CD45 isoforms and CCR7, which distinguish T cell subsets in several species, were used for initial characterization of CD8 T cells from different tissues of Babraham pigs. BAL, which contains ∼80% macrophages as in other species, is of particular interest as it contains airway T cells that are at the frontline of protection against respiratory pathogens (*33*). Among the lymphocytes in BAL ∼20% were CD4 T cells, ∼20% CD8 and a slightly higher proportion γδ T cells (*22*).

Because porcine T helper cells express CD8α when activated (*28, 34*), we used an antibody to CD8β combined with CD45RA and CCR7 antibodies to identify and characterize CD8 T cells. Four populations were apparent (**Fig. 1A**) in blood, spleen and TBLN, which appear to correspond to those defined in humans as naïve (CD45RA^+^CCR7^+^), central memory (TCM) (CD45RA^-^CCR7^+^), effector memory (TEM) (CD45RA^-^CCR7^-^) and terminally differentiated effector cells (TDE) (CD45RA^+^CCR7^-^) (*35*). The mucosal tissues (lung and BAL) contained higher proportions of cells with TEM and TDE phenotypes. To confirm this differentiation scheme for CD8 T cells, we examined expression of CD27, a marker of less differentiated cells and perforin, expressed by effector CD8 T cells. These experiments indicated that the proposed naïve and TCM cells in all tissues expressed CD27 but little or no perforin, while TEM and TDE cells were heterogeneous, expressing one or the other marker or neither, as in humans (**Supplementary Fig. 1**) (*35–37*). These data support the identification of naïve and TCM cells and indicate that TEM and TDE are more differentiated cells with effector function. This phenotypic differentiation was further confirmed by examining cytokine production of naïve, TCM, TEM and TDE CD8 T cells sorted from peripheral blood mononuclear cells (PBMC) following polyclonal stimulation with PMA/ionomycin. A high proportion of TEM produced IFNγ and TNF while lower proportions of TDE and TCM and few naïve cells did so (**Fig. 1B, C, D**). Naïve and TCM cells secreted mainly TNF (4.4% and 8.7% respectively) while many TEM were double producers (IFNγ^+^TNF^+^ 18.9%). TDE produced predominantly IFNγ (8%) with a smaller proportion secreting both IFNγ and TNF (4.9%).

**Figure 1.**
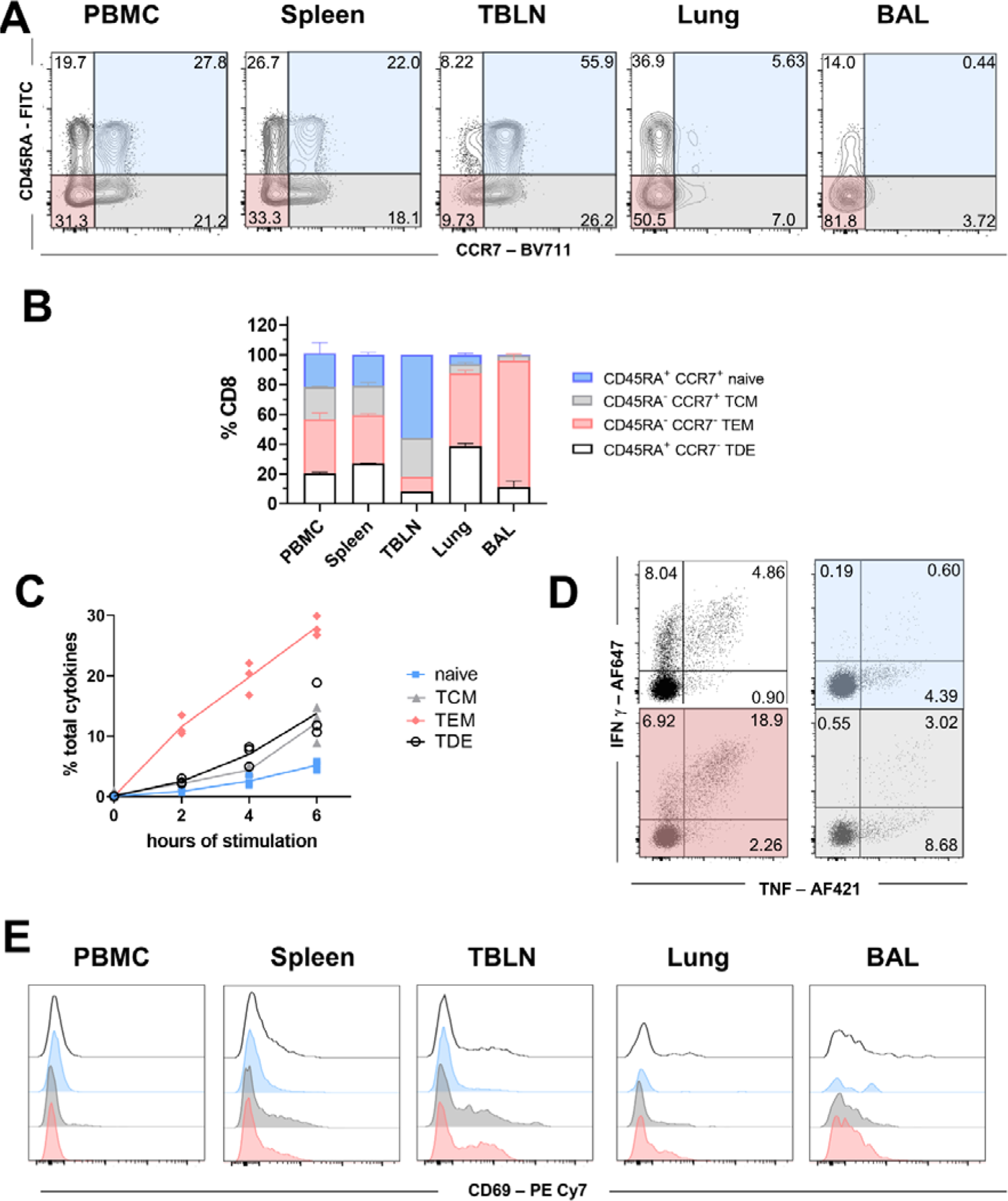
Phenotype of porcine CD8 T cells in tissues and cytokine secretion after stimulation. **(A)** Expression of CD45RA and CCR7 by CD8 T cells isolated from the indicated tissues of naïve Babraham pigs. Quadrants show the proportion of each population. **(B)** Mean frequency (± SD) of TDE (CD45RA^+^, CCR7^-^), naïve (CD45RA^+^, CCR7^+^), TEM (CD45RA^-^, CCR7^+^) and TCM (CD45RA^-^, CCR7^-^) in CD8 T cells from 3 animals. **(C)** CD8+ T cells from PBMC were sorted according to their expression of CD45RA and CCR7. The sorted cells were stimulated with PMA and Ionomycin for 0, 2, 4 and 6 hours and TNF and IFNg secretion measured by intracellular cytokine staining. Each symbol represent one animal, this experiment was repeat twice**. (D)** Representative FACS plot showing the secretion of IFNg and TNF after 6 hours stimulation in TDE (white panel), naïve (blue panel), TCM (grey panel) and TEM (red). Mean proportion of IFNg single (top left), double producer (top right) and single TNF^+^ (bottom right) T cells are reported. **(E)** CD69 expression in terminally differentiated effector (TDE, white), naïve (blue), central memory (TCM, grey) and effector memory (TEM, red) in CD8 T cells in the tissues analyzed.

We have previously shown that BAL and a large proportion of lung T cells are not stained by intravenous anti-CD3 antibody (*23, 25*), indicating that these populations are TRM. It is striking that BAL T cells are almost exclusively highly differentiated. Staining with CD69, a marker of activation and tissue residency (*38, 39*) showed that, as in humans, CD69 was absent or minimally expressed on blood CD8 T cells, while the highest expression was found on TCM and TEM in TBLN and on T cells in BAL. CD8 TDE in the lungs expressed lower levels of CD69 (**Fig. 1E**).

These data demonstrated that in pigs, CD69 expression is tissue dependent on activated/memory CD8 T cells. BAL CD8 T cells are predominantly TEM phenotype with high expression of CD69 and lack of staining by intravenous CD3 antibody, indicating that they are TRM. TEM are heterogenous in expression of CD27 and perforin and produce high levels of cytokines on stimulation.

### Kinetics and phenotype of influenza-specific CD8 T cells

Having established the phenotype of CD8 T cells from unimmunized pigs, we wished to define the temporal dynamics of the immune response to IAV. Inbred Babraham pigs were infected with H1N1pdm09 in four experiments. One pig was culled on days 1-7, 9 and 11 post infection (DPI) in two experiments, three more at 6, 7, 13, 14, 20 and 21 DPI and a further four at 21, 42 and 63 DPI in additional experiments (*22*) (**Fig. 2A**). We determined the proportions of influenza NP-specific CD8 T cells binding to tetramers carrying the DFE epitope and tetramers of the newly identified AAV and VAY epitopes (**Supplementary Fig. 2**) in various tissues over time and modelled it, starting at 6 DPI when a cellular response is first detectable (*22*). The results and curves fitted to the data are shown in **Fig. 2B** and indicated that the peak proportion of CD8^+^ T cells specific for each tetramer and the timing of the peak differed amongst tissues and between tetramers. The highest response for all tetramers, expressed as a percentage or absolute count, was found in the BAL, followed by lung and local lymph nodes, (**Fig. 2B** and **Supplementary Fig. 3**). Interestingly, the modelled response in PBMC peaked earlier (16.8 DPI for AAV, 3.4 DPI for DFE and 9.1 DPI for VAY) compared to BAL, lung or nasal turbinates (NT) (28.0 DPI, 11.2 DPI and 11.1 DPI respectively in BAL) (**Table 1**), in accordance with the idea that cells generated in lymph nodes traffic to local tissues via the blood. AAV responses peaked 3 to 17 days later than the peak of DFE or VAY responses in different tissues, while DFE and VAY shared similar kinetics in most tissues except PBMC, where DFE peaked earlier (**Table 1** and **Fig. 2B**). The magnitude of AAV responses was greater than those to VAY and DFE in all tissues, while DFE was higher than VAY in all tissues except for TBLN (**Table 1** and **Fig. 2B**). At the later time points of 21, 42 and 63 DPI we also examined tracheal CD8 T cells and here too the AAV response was higher than that to DFE and VAY at 42 DPI (p=0.03), with DFE being significantly higher than VAY (p=0.03). AAV remained significantly higher than VAY at 63 DPI (p=0.03) (**Supplementary Fig. 4A and B**).

**Figure 2.**
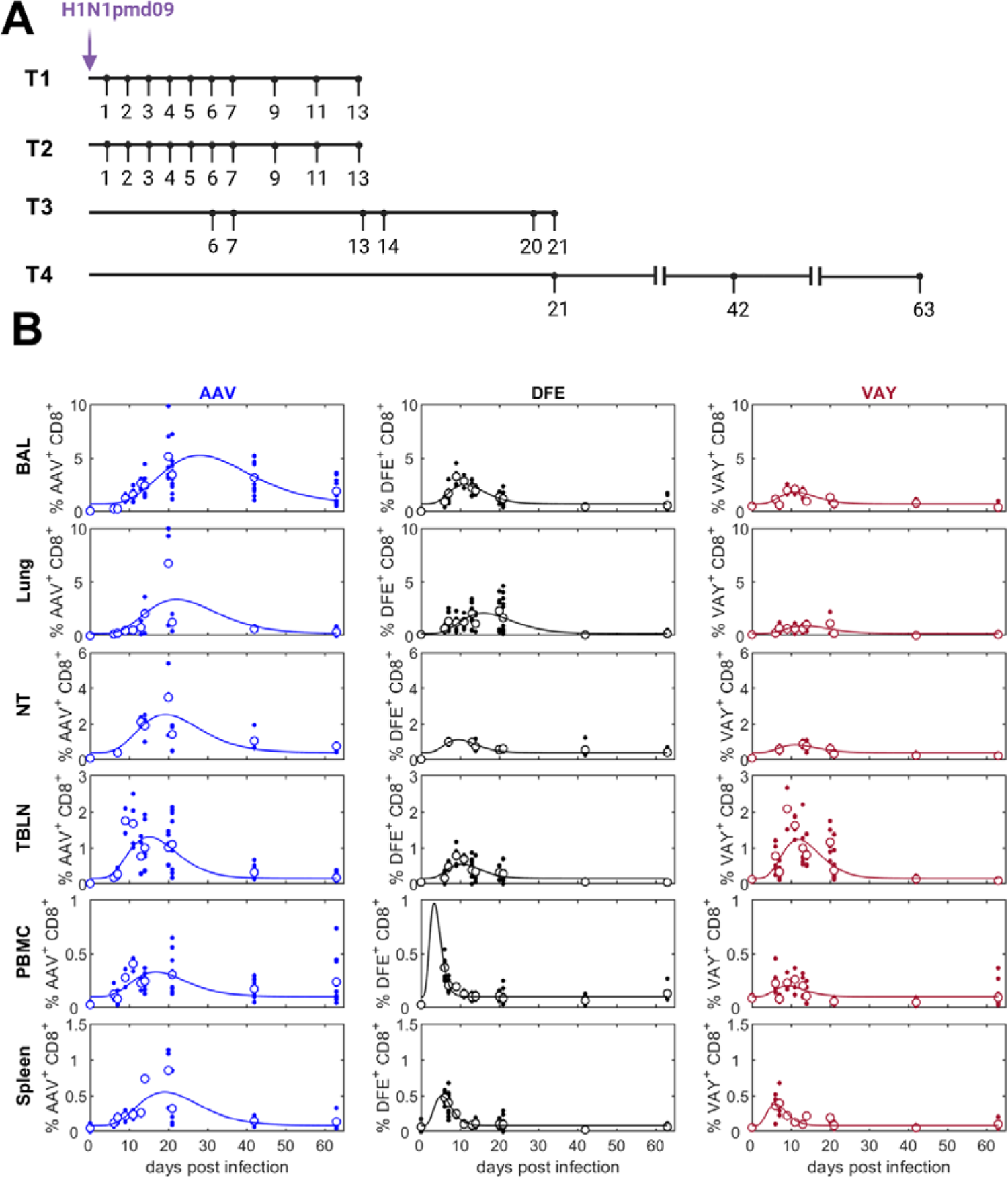
Experimental design and tetramer distribution in tissues. **(A)** Babraham pigs were infected with H1N1pdm09 intranasally and culled on the indicated days post infection. Broncho-alveolar lavage (BAL), lung, tracheo-bronchial lymph nodes (TBLN), peripheral blood mononuclear cells (PBMC) and spleen were collected at all time points while nasal turbinates (NT) were isolated only in experiments T3 and 4. **(B)** Proportion (%) of CD8^+^ T cells specific for AAV (left), DFE (middle) or VAY (right) in different tissues. In each plot the solid line is the fitted curve describing the dynamics, the large open circles are the mean % at each time point and the small filled circles are the observed % for individual pigs at each time point.

**Table 1.** Estimated parameters describing the changes over time in the proportion of CD8^+^ T cells specific for each tetramer in different tissues.

| parameter | tissue* | Tetramer |  |  |
| --- | --- | --- | --- | --- |
|  |  | AAV | DFE | VAY |
| peak proportion (%) | BAL | 5.23 | 2.52 | 1.80 |
|  | Lung | 3.37 | 2.10 | 0.90 |
|  | NT | 2.53 | 1.12 | 0.83 |
|  | TBLN | 1.31 | 0.55 | 1.23 |
|  | PBMC | 0.33 | 0.97 | 0.20 |
|  | Spleen | 0.55 | 0.50 | 0.38 |
| time of peak proportion<br>(days post infection) | BAL | 28.0 | 11.2 | 11.1 |
|  | Lung | 22.1 | 15.9 | 13.6 |
|  | NT | 19.3 | 9.5 | 11.1 |
|  | TBLN | 15.1 | 10.3 | 11.9 |
|  | PBMC | 16.8 | 3.4 | 9.1 |
|  | Spleen | 19.1 | 5.3 | 6.0 |
| baseline proportion (%)† | BAL | 0.71 |  |  |
|  | Lung | 0.22 |  |  |
|  | NT | 0.41 |  |  |
|  | TBLN | 0.16 |  |  |
|  | PBMC | 0.10 |  |  |
|  | Spleen | 0.09 |  |  |

\* BAL - broncho-alveolar lavage; NT - nasal turbinate; TBLN - tracheo-bronchial lymph node
† baseline proportion is the same for each of the tetramers

We next modelled changes in the proportions of the different tetramer^+^ cells in each tissue. BAL, lung and spleen showed similar changes in proportions of the different tetramer^+^ populations. Initially AAV was lower than VAY and DFE in all tissues but by 30 DPI AAV was dominant (60%), while in most tissues DFE and VAY declined (∼20%) (**Fig. 3A**). However, in NT the frequency of cells specific for DFE remained constant (25%), while that for VAY decreased and that for AAV increased. In PBMC the proportions changed only gradually (**Fig. 3A**), though the small number of PBMC data points indicate that this observation should be interpreted with caution.

**Figure 3.**
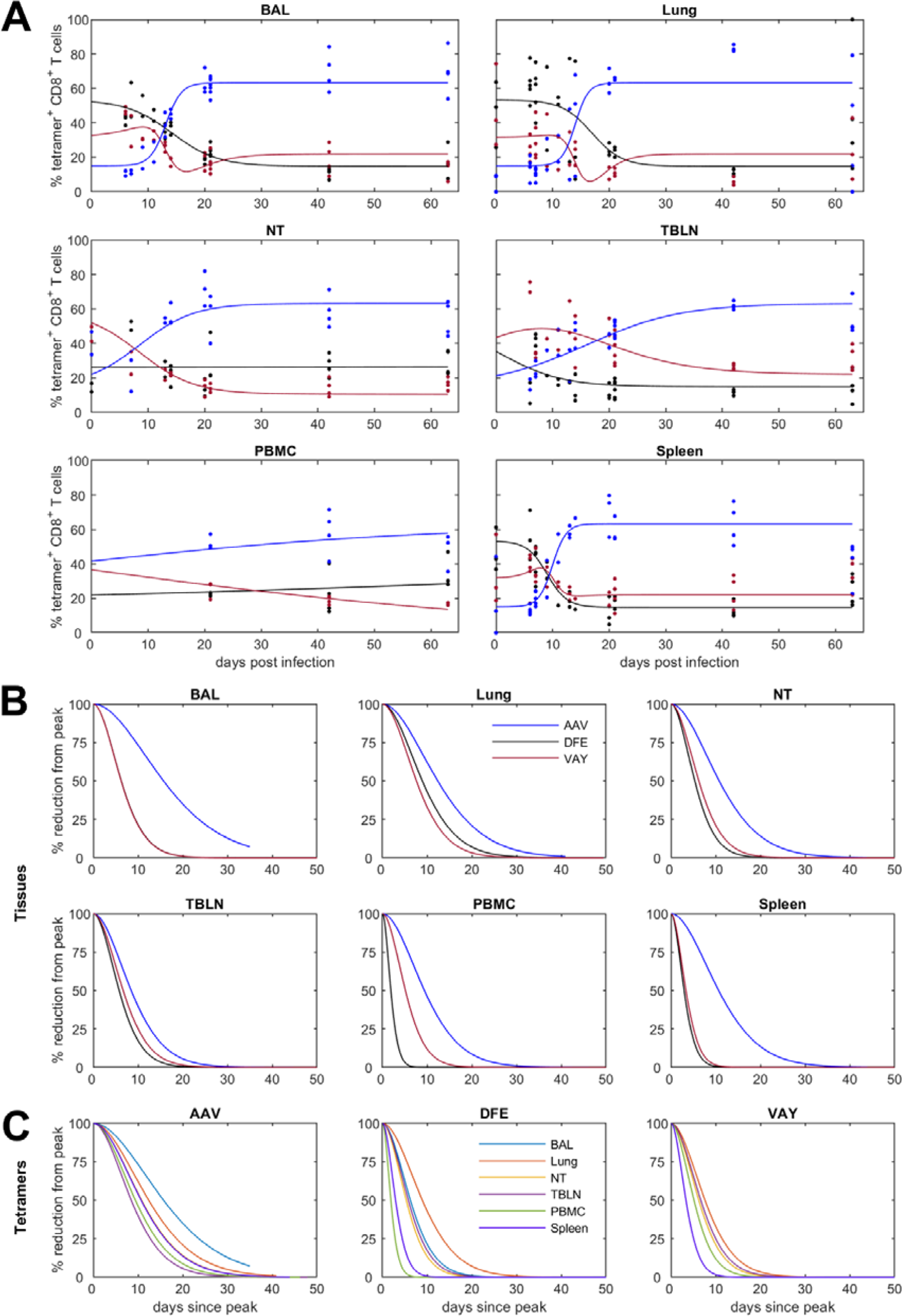
Changes in the proportion of tetramer^+^ CD8 T-cells in different tissues and estimated decay (% reduction from peak) over time. **(A)** Relative proportion (%) of CD8^+^ T cells specific for each tetramer (AAV – blue; DFE - black; VAY - red) in the indicated tissues. In each plot the solid line is the fitted curve describing the dynamics and the points are the observed proportions for individual pigs at each time point. **(B)** Decay from the peak of response within the indicated tissue. **(C)** Decay of tetramer^+^ CD8 T-cells in different tissues (as indicated). Note: DFE and VAY have the same dynamics in BAL, VAY has the same dynamics in NT and PBMC and AAV has the same dynamics in NT and spleen so these decay curves overlap.

Influenza specific T cells in the nasal mucosa of mice have been shown to be longer lived and decline less rapidly than those present in the harsh environment of the lung (*40*). Using a mathematical model, we investigated the decline of tetramer^+^ T cells in all tissues after the peak response (**Fig. 3B**). The decay in the proportion of cells specific for AAV was slower than that for cells specific for either DFE or VAY in all tissues. Furthermore, the proportion of cells specific for AAV decayed most slowly in BAL.

Collectively, our results shows that the frequency of different tetramer^+^ T cells varies between tissues, with the highest frequency in BAL. AAV tetramer^+^ cells are dominant at later time points. In general, responses in PBMC peaked earlier compared to local tissues but waned more rapidly and did not reflect events in mucosal tissues. We did not observe a more rapid decay in lung and BAL compared to NT, as previously reported in mice (*40*).

### BAL T cells maintain a stable phenotype but transcription alters over time

We next analyzed the phenotype of the NP-specific T cells present locally (in BAL and TBLN) and systemically (PBMC) (**Fig. 4A**). The majority of tetramer^+^ cells were TCM or TEM throughout the time course in all tissues analyzed. There was a slow but steady increase in the proportion of TCM with time in PBMC, reflecting the phenotype found most abundantly in TBLN (**Fig. 4B**). On average, more than 80% of influenza-specific T cells in local lymph nodes were TCM, while only a small proportion (8%) were TEM (**Fig. 4A and B**). In contrast, in BAL the majority of cells are TEM (78.5%), with only a small number of TCM (20.3% at 21 DPI) (**Fig. 4A and B**). Changes in TDE and naïve cells are not shown as the numbers of these cells were too small for reliable analysis.

**Figure 4.**
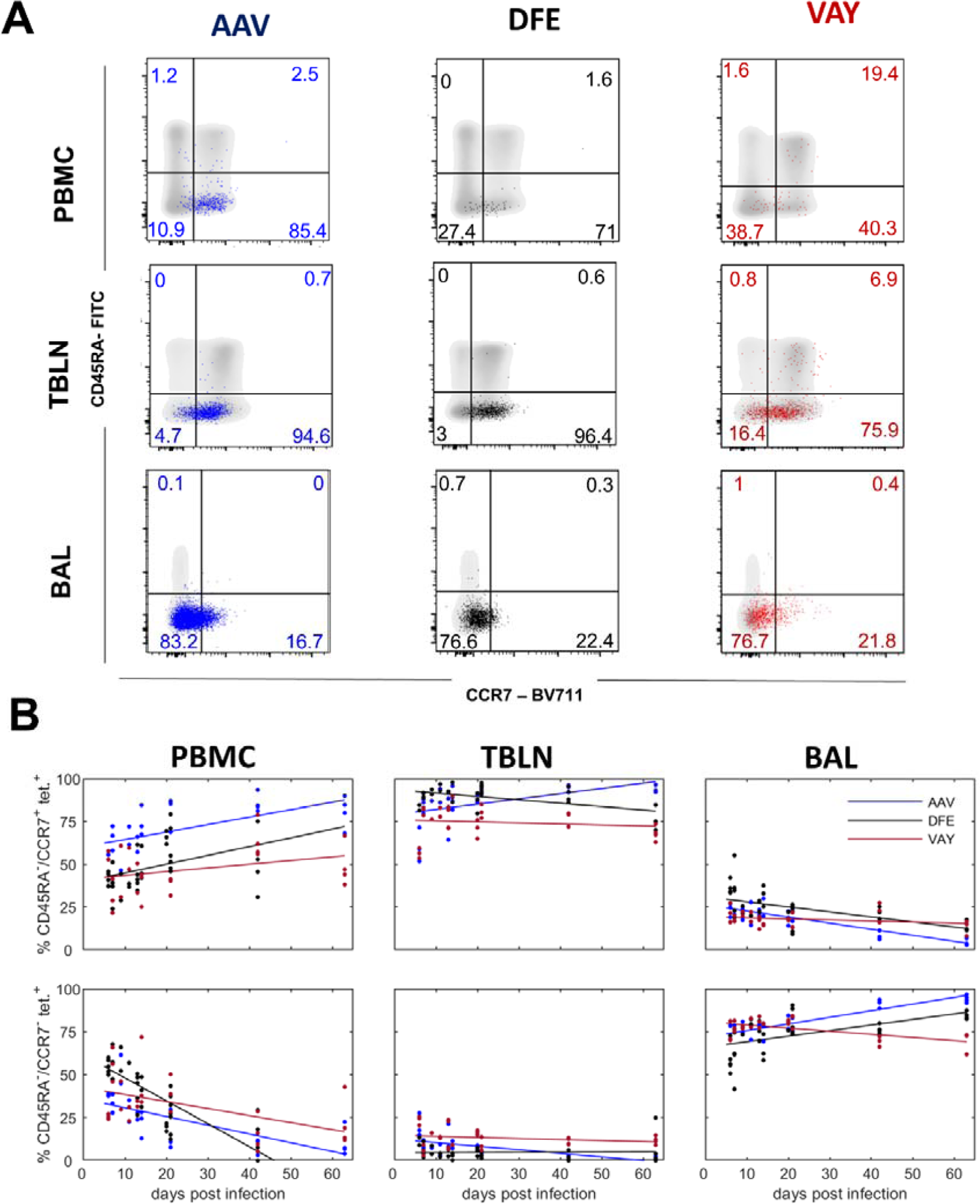
Phenotype of influenza-specific CD8 T cells in tissues over time. **(A)** Expression of CD45RA and CCR7 by AAV (left, blue), DFE (centre, black) and VAY (right, red) tetramer^+^ (coloured dots) and total CD8 T cells (in grey) isolated at 21 DPI from PBMC (top), TBLN (middle) BAL (bottom), representative plots for one individual. **(B)** Proportion (%) of CD8^+^ T cells staining with AAV (blue), DFE (black) and VAY (red) tetramer in different tissues. T cell populations are TEM (CD45RA^-^/CCR7^+^; top row) and TCM (CD45RA^-^/CCR7^-^; bottom row) in PBMC (left), TBLN (center) and BAL (right). In each plot the solid line is the fitted trend and the points are the observed proportions for individual pigs at each time point.

We used RNA sequencing (RNA-seq) to compare the transcriptome of CD8 T cells specific for the previously defined DFE NP epitope (*32*). DFE^+^ T cells were isolated from BAL by cell sorting at different time points (7, 21 and 63 DPI) (**Fig. 5A**). Differential gene expression analysis was applied to compare these three groups. 4,666 genes were expressed at significantly different levels (p_adj_ ≤ value 0.05 and |log_2_ fold change| > 1) at 7 DPI versus (vs) 21 DPI while 1,198 were differentially expressed in 7 DPI vs 63 DPI and only 560 in 21 DPI vs 63 DPI (**Fig. 5B**). At 7 DPI several upregulated genes were involved in cell growth, movement (*Igfbp2*) and proliferation (*Ctla4,Kif11, Kif18a, Shmt1*) while at 21 DPI genes linked with T cell activation (*Tagap, IL2ra, Csf2, Dgkg*) and adhesion (*L1cam, Cass4*) were highly expressed (**Fig. 5C**). Interestingly, comparison of 7 DPI and 63 DPI revealed differential expression of genes involved in metabolism (*Jazf1, Atp8b4, Igf2bp3,*), transcription factors (*Litaf*), T cell development and proliferation (*Ccnd3, Shcbp1*). Upregulated pathways at 21 DPI, compared to 7 DPI, were linked to the control of Th1/Th2 differentiation, cytokine secretion, antigen processing and presentation (**Fig. 5D, Table S1**). The TGFβ signalling pathway, known to be involved in mucosal residency, was also upregulated at 21 DPI.

**Figure 5.**
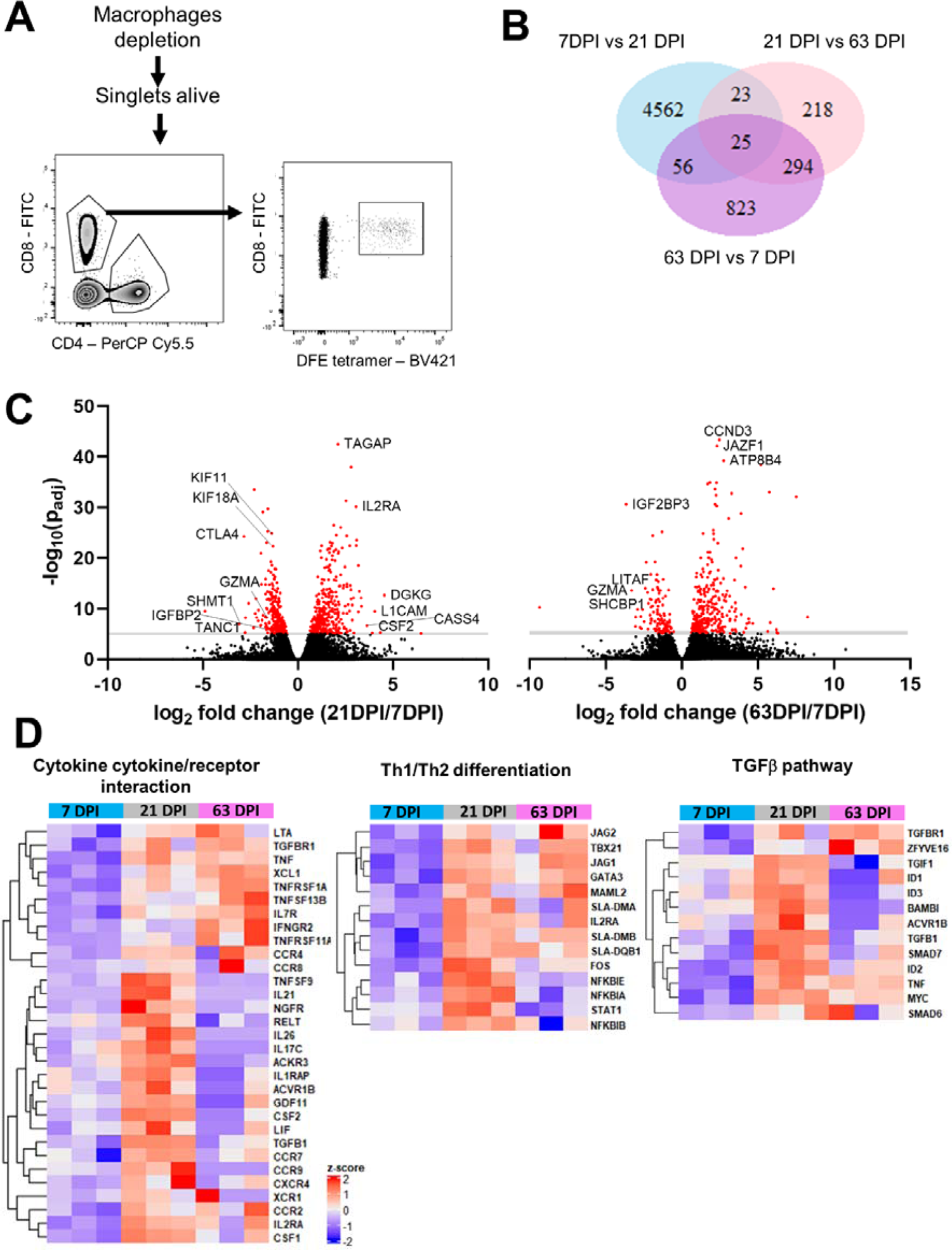
Transcriptional profile of DFE-specific T cells in BAL at 7, 21 and 63 DPI. **(A)** Sorting strategy for the isolation of DFE^+^ T cells for RNA-seq. **(B)** Venn diagram shows the number of significant differentially expressed genes (p_adj_ value<0.05, and |log_2_ fold change|> 1) between 7DPI versus (vs) 21 DPI, 21DPI vs 63DPI and 63DPI vs 7DPI samples. **(C)** Volcano plot showing upregulated genes in 21DPI vs 7DPI and 63DPI vs 21DPI comparison. **(D)** Heatmap of selected genes from KEGG Pathway analysis related to cytokine production, T cell differentiation and TGFβ pathway (enrichment score of 0.76, 0.88 and 0.83 respectively)

We next examined the presence of key gene expression features of TRM, previously identified in the human lung (*12, 41, 42*) (**Supplementary Fig. 5**). BAL cells from 63 DPI upregulated a gene related to integrins (*Itga1*), the TRM transcription regulator gene *Znf683* and downregulated genes involved in migration (*Sell, S1pr1*) as in humans. In addition, at the earlier timepoint of 7 DPI DFE^+^ T cells expressed more cytotoxicity related genes (*GzmA, GzmH, Prf1* and *Ccl5*) while starting from 21 DPI genes involved in cytokine signalling and secretion were upregulated (*Ifng, Tnf, Il13, Tgfb1* and *Tnfsf13b*) (**Supplementary Fig. 5**). Interestingly, *CD69* expression changed with time with its peak at 21 DPI, as did CD103 gene expression (*Itgae*) with a peak at 7 DPI.

Despite the similar phenotype, these data suggests that gene expression in CD8 T cells at the site of infection changes over time, with genes involved in proliferation and migration upregulated at 7 DPI while cytokine related pathways are upregulated at 21 DPI.

### BAL tetramer^+^ cells are a stable highly differentiated population

RNA-seq analysis revealed changes in CD69 gene expression with time, we therefore studied CD69 protein expression and modelled these changes in BAL, TBLN and PBMC (**Fig. 6A**). CD69 expression decayed only slightly with time in BAL for all tetramers and also for AAV and VAY labelled T cells in TBLN, while it decayed in TBLN DFE^+^ T cells at a higher rate (0.022/day for DFE, 0.003/day VAY and 0.001/day AAV) (**Table 2 and Fig. 6A**). As reported above, PBMC expressed minimal CD69.

**Figure 6.**
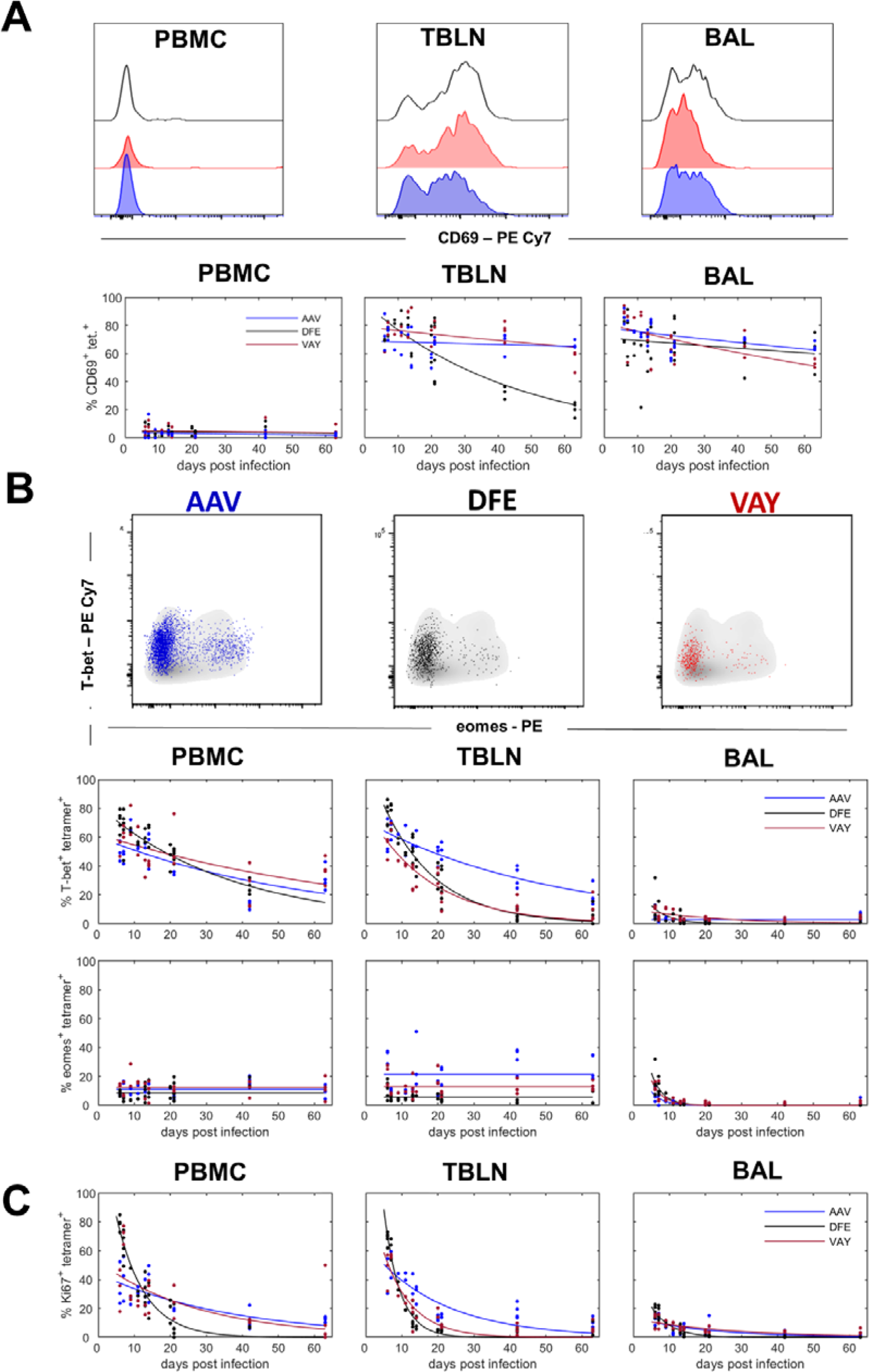
Activation state and transcription factors expression in tetramer^+^ CD8 T cells. **(A)** Top: Histograms show the expression of CD69 by DFE (black), VAY (red) and AAV (blue) tetramer^+^ T cells in PBMC (left), TBLN (centre) and BAL (left) at 21 DPI. Bottom: Frequencies of tetramer+ cells expressing CD69 in PBMC, TBLN and BAL. **(B)** Top: Expression of T-Bet and Eomes by AAV (left, blue), DFE (centre, black) and VAY (right, red) tetramer^+^ (coloured dots) and total CD8 T cells (in grey) isolated at 21 DPI from TBLN, representative plots for one individual. Bottom: Frequencies of tetramer+ cells expressing T-bet and Eomes in PBMC, TBLN and BAL. **(C)** Proportion of AAV, DFE and VAY tetramer ^+^ expressing Ki67 in PBMC, TBLN and BAL. In each plot the solid line is the fitted curve describing the dynamics and the points are the observed proportions for individual pigs at each time point

**Table 2.** Estimated decay rates (*d*; /day) in the proportion of tetramer specific T cells expressing different markers across tissues.

| marker | tissue* | Tetramer |  |  |
| --- | --- | --- | --- | --- |
|  |  | AAV | DFE | VAY |
| Ki67 | PBMC | 0.027 | 0.128 | 0.036 |
|  | TBLN | 0.049 | 0.226 | 0.127 |
|  | BAL | 0.050 | 0.186 | 0.038 |
| T-bet | PBMC | 0.017 | 0.027 | 0.013 |
|  | TBLN | 0.019 | 0.067 | 0.057 |
|  | BAL | 0 | 0.196 | 0.044 |
| EOMES† | PBMC | 0 |  |  |
|  | TBLN | 0 |  |  |
|  | BAL | 0.33 |  |  |
| CD69 | PBMC | 0.012 | 0.006 | 0.006 |
|  | TBLN | 0.001 | 0.022 | 0.003 |
|  | BAL | 0.004 | 0.003 | 0.007 |

\* BAL - broncho-alveolar lavage; TBLN - tracheo-bronchial lymph node
† decay rate is the same for each of the tetramers

Pathway analyses revealed upregulation of Th1/Th2 differentiation related genes in BAL T cells at 21 DPI compared to 7 DPI. We validated these findings by analyzing and modelling the expression of the transcription factor T-bet, involved in Th1 differentiation and homing to inflammatory sites, and Eomesodermin (Eomes), involved in induction of memory and effector T-cell differentiation (*43*) (**Fig. 6B**). Eomes was poorly expressed in BAL, only detectable at early time points and decayed rapidly in all tetramer^+^ cells (0.33/day decay rate) (**Table 2**). The highest expression of this transcription factor was found in AAV^+^ T cells in TBLN (mean of 20.6%) followed by VAY and DFE while similar expression was present in PBMC for all tetramers, with no decay (**Fig. 6B** and **Table 2**). In contrast, T-bet expression differed greatly among tissues and tetramers. T-bet was highly expressed in TBLN and PBMC, where it decayed more slowly than in TBLN for all tetramers. In TBLN, T-bet decreased more gradually in AAV than DFE or VAY responding T cells (**Fig. 6 B** and **Table 2**). There was low expression of T-bet at early times in BAL, which declined rapidly and was undetectable at later time points in all tetramers (**Table 2**).

These data suggested that BAL TRM may have already switched off Eomes and T-bet protein expression and are no longer undergoing active Th1/Th2 differentiation. We therefore analysed Ki67 expression as a proxy for cell proliferation which is normally linked to differentiation. Whereas high frequencies of Ki67^+^ tetramer binding cells were found in PBMC and TBLN at early time points, only 14% of BAL cells were Ki67^+^ at 6 DPI and Ki67 expression was barely detectable at 63 DPI (1.8%) (**Fig 6 C**).

The RNA-seq data indicates changes in expression of many genes related to cytokine production over time. Lymphocytes isolated from PBMC, TBLN and BAL were therefore stimulated with H1N1pdm09 and the production of IFNγ, TNF and IL-2 by tetramer binding cells was assayed by intracellular cytokine staining. We compared the responses of the dominant responding AAV population with DFE cells which decline more rapidly. Despite high expression of T-bet, PBMC tetramer^+^ cells produced a limited amount of IFNγ (5.9% in DFE^+^, 4,3% AAV^+^ at 7 DPI) which was almost undetectable after 21 DPI (**Fig. 7A and B**). Similar kinetics were found in TBLN, with IFNγ and IFNγ/TNF co-producing cells being the most abundant. The highest responses were in BAL, with consistent production of cytokines (predominately IFNγ and TNF) even at 63 DPI. No significant difference was observed between DFE^+^ and AAV^+^ populations, but there was a trend toward a higher proportion of triple producers (INFγ^+^TNF^+^IL-2^+^) at later time points in AAV^+^ cells compared to DFE^+^ (**Fig. 7 B**).

**Figure 7.**
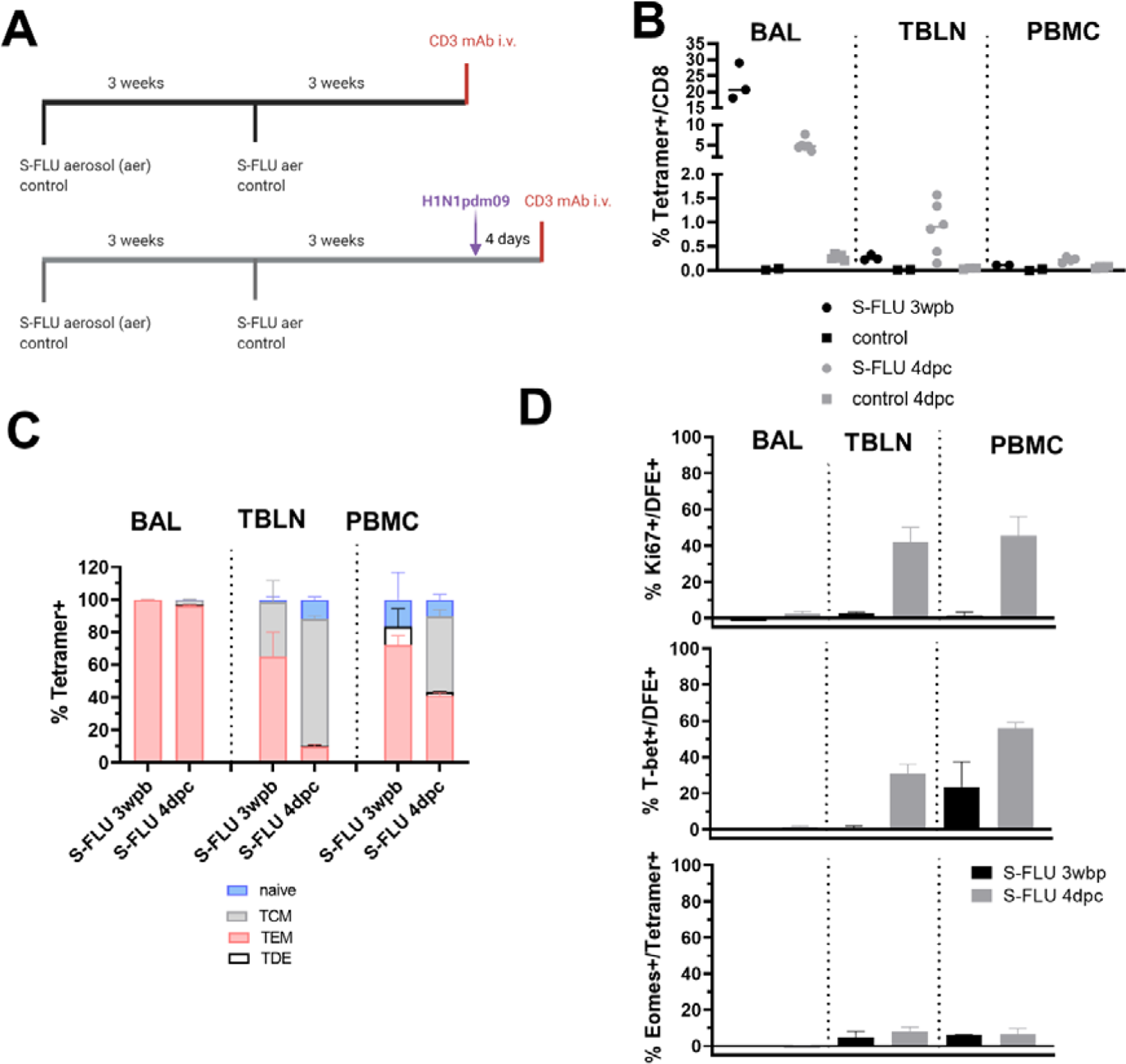
Cytokine secretion after *ex vivo* virus stimulation of AAV and DFE tetramer^+^ cells. **(A)** Lymphocytes isolated at different time points from blood, TBLN and BAL were stimulated with H1N1pdm09 MOI=1. Following 18 hours of stimulation, cells were labelled using tetramers and cytokines quantified using intracellular cytokine staining. Representative plots of AAV^+^ (on left) and DFE^+^ (right) T cells secreting IFNg, TNF and IL-2 cytokines, from lymphocytes isolated 14 DPI. **(B)** Mean frequency (± SEM) of tetramer^+^ cells secreting IFNg, TNF and IL-2 from PBMC (top panels), TBLN (in the middle) and BAL (bottom). The right Y axes shows the mean frequency of AAV^+^ (in blue, left panels) and DFE^+^ (in black, right panels) within CD8 T cells. Data shown are mean of 3 / 4 individuals per timepoint. Two-way ANOVA was used for comparison of each cytokine population between DFE^+^ and AAV^+^ cells.

In conclusion, despite differences in transcription of CD69 and T cell differentiation genes, we did not find corresponding differences in the protein level expression of CD69, T-bet and Eomes in BAL, while TBLN and PBMC expressed T-bet and Ki67 at early time points and EOMES up to 63 DPI. *Ex vivo* stimulation with H1N1pdm09 resulted in IFNγ and TNF cytokine secretion at 7 DPI, while upregulation of related genes was highest at 21 DPI. These data suggest that BAL TRM are a stable largely non-dividing population able to produce abundant cytokines for at least 63 DPI.

### Aerosol immunization generates a powerful CD8 response with a similar phenotype to influenza infection

To investigate whether immunization elicits similar responses to natural infection we administered S-FLU, a single cycle influenza vaccine, by aerosol to Babraham pigs and boosted them 3 weeks later. In a first experiment three pigs were culled 3 weeks post boost (WPB) while in a second, S-FLU immunized pigs (n=6) were challenged with H1N1pdm09 and culled four days later (**Fig. 8A**). Control pigs were left untreated in the first or were challenged without prior immunization in the second experiment. Anti-porcine CD3 mAb was administered intravenously (i.v.) 10 minutes prior to sacrifice to distinguish between tissue resident (CD3 i.v.^-^) and circulating T cells (CD3 i.v.^+^) (**Fig. 8A and Supplementary Fig. 6**). We enumerated S-FLU specific cells using the DFE tetramer, as this epitope is conserved in the vaccine and challenge viruses. In BAL, 22.6% of CD8 T cells were DFE^+^ at 3 WPB but only 5% 4 days post challenge (DPC) (**Fig. 8 B**). Conversely, there were higher numbers of TBLN and PBMC DFE^+^ cells 4 DPC compared to 3 WPB (0.8% vs 0.3% in TBLN and 0.2% vs 0.1% in PBMC) (**Fig. 8 B**). As we have shown before, BAL and TBLN cells were completely inaccessible to blood (CD3 i.v.^-^) while most spleen cells were labelled with the infused antibody (**Supplementary Fig. 6**), confirming that BAL cells are truly tissue resident and that lymph node cells are also outside the blood stream (*23, 25*).

We then used CD45RA and CCR7 to study differences in phenotype. Three weeks after S-FLU immunization, BAL cells were almost exclusively TEM as after natural infection and this did not change after challenge (**Fig. 8C**). Following challenge however, there was a rapid increase in the proportion of TCM in TBLN and PBMC (from 33% at 3 WPB to 77% at 4 DPC in TBLN and 0% to 46% in PBMC).

In the earlier H1N1pdm09 infection experiment, few BAL CD8 were proliferating. We therefore examined whether DFE^+^ T cells expressed Ki67 early after immunization and challenge with live virus. Ki67 was absent at 3 WPB in all tissues, and only marginally expressed at 4 DPC in BAL (2.7%). In contrast 42.2% of DFE^+^ TBLN cells and 45.4% of DFE^+^ PBMC expressed Ki67 after challenge (**Fig. 8D and E**). As during influenza infection, in S-FLU immunized pigs DFE^+^ BAL cells lacked T-bet and EOMES while these transcription factors were expressed in TBLN. T-bet expression reached 30.6% 4 DPC (from 1% at 3 WPB) while EOMES was only slightly upregulated (4.7% at 3 WPB and 7.9% at 4 DPC). Tetramer^+^ T cells in PBMC expressed T-bet at 3 WPB (23.5%) and it increased after challenge (55.9%) with no changes in EOMES expression (6.2% 3 WPB and 6.7% 4 DPC) (**Fig. 8D**).

Taken together these results suggest that S-FLU immunization by aerosol elicited a stronger T cell response in BAL compared to influenza infection but the responding T cells had a similar differentiated, non-proliferating phenotype. Perhaps surprisingly the proportion of DFE tetramer^+^ cells in BAL decreased four days after challenge with infectious virus.

## Discussion

In this study we have described in detail for the first time porcine influenza-specific CD8 T cells in lymphoid and non-lymphoid tissues and shown that antibodies to CD45RA and CCR7 identify four subsets: naïve, TCM, TEM and TDE. The identity of these subsets was substantiated by additional staining for CD27 and perforin, and by assessing their ability to secrete effector cytokines. We have also shown that while all four CD8 T cell subsets are well represented in PBMC, spleen and TBLN, the subsets show a very different distribution in lung tissue and BAL. In lung tissue the majority of CD8 cells are TDE and TEM as has been previously reported in humans (*41*), while in BAL TEMs prevail (∼80%). We have already shown that the majority of TBLN, BAL and, to a lesser extent, lung T cells are inaccessible to intravenous anti-CD3 antibody, it was therefore of interest to examine the expression of CD69, designated as a marker of activation and tissue residence, on these cells. As in other species, we found that CD69 is minimally expressed on blood T cells although upregulated if PBMC are activated with a mitogen (data not shown), but in all the other tissues examined, some cells express this antigen. In BAL, CD69 is expressed on a high proportion of CD8 T cells, but lower levels of CD69^+^ cells are found in the lung. In TBLN all the CD8 subsets show clear CD69^-^ and CD69^+^ populations and in the spleen we could also detect CD69^+^ cells, perhaps reflecting that a proportion of cells in these organs may have been recently activated. In summary, a high proportion of NP specific T cells in TBLN expresses CD69 from 6 days after influenza challenge until 63 DPI (**Fig. 6A**).

Due to the identification of two new CD8 epitopes in influenza NP, we were able to compare changes in the distribution, phenotype and gene expression of different antigen specific cells during influenza infection (*32*). We tracked CD8 T cells responding to three NP epitopes in blood and tissues from 6 to 63 days post infection. Intriguingly, the modelled kinetics of the responses to the three epitopes are very different, with DFE tetramer numbers showing a very early peak in PBMC and spleen but peaking in all other tissues at similar times to VAY. The response to the third epitope AAV appears later, but by 20-30 DPI it is dominant in all tissues and persists for longer than the response to the other two epitopes.

Others have reported marked differences in the hierarchy of the responses to immunodominant epitopes of influenza in mice (*44, 45*). Pizzolla and colleagues have shown that T cells specific for immunodominant epitopes in NP, PA and PB proteins of influenza in the nasal mucosa shared a similar hierarchy with systemic responses while in the lung all immunodominant epitopes were equally represented (*40*). In our model, despite an initial peak of DFE tetramer stained T cells, at the memory stage, the AAV epitope dominates in all tissues, with no marked difference between the organs. Furthermore, the response of tetramer positive cells does not decline more rapidly in the lung or BAL than blood or spleen, although in mice both of these have been postulated to be hostile environments inducing transcriptional and epigenetic changes that promote apoptosis (*46*). Further studies are required to understand why AAV responses are immunodominant compared to DFE and VAY. Immunodominance may be linked to peptide-MHC affinity and it has also been reported that early IFNγ production can provide an advantage to a given antigen specific population (*47, 48*). However, we did not observe significant differences in cytokine production of AAV^+^ and DFE^+^ CD8 T cells (**Fig. 7**).

Mechanisms by which TRM populations persist have been investigated by several authors in mice with sometimes contradictory results, so that it remains unclear whether some or all respiratory tract TRM populations are maintained by recruitment of circulating cells. While some experiments suggest that antigen encounter in the lung environment is important for TRM establishment and maintenance (*40, 46, 49, 50*), it has been reported that lung TRM may develop independently of pulmonary antigen encounter under specific inflammatory conditions (*51*). Other results indicate that TEM but not TCM can be recruited to the lungs and since relative numbers of TEM in the circulation decrease with time after antigen exposure while TCM increase, as observed in PBMC (**Fig. 4B**) this may explain why recruitment of TRM declines with time (*51*). These data are in accord with parabiosis experiments in mice since if the parabiosis is carried out early after immunization, (activated) antigen-specific cells from the immunized animal enter the lungs of the parabiotic partner but not if the parabiosis is performed later (*50*).

Our data indicates that following influenza infections an airway population of TRM, recoverable by BAL, is established with a peak at 20-30 DPI of predominantly AAV-specific cells. We hypothesize that the late dominance of AAV may be because cells with this specificity continue to divide for longer than those specific for DFE or VAY, with continuing generation of cells with a TEM phenotype that can enter the lungs. Early on, BAL TRM cells express genes characteristic of T cell activation and 5-22% express Ki67, though Ki67 expression is much more prominent in tetramer binding cells in TBLN and PBMC, suggesting that the bulk of the BAL population arises by cell division prior to entry into the airways. BAL Ki67 expression declines over time and, since the tetramer binding population also declines, cell division clearly does not maintain the population. This finding is in line with recent studies on SARS-CoV-2 infection, revealing that despite a high number of lung-resident T cells, these lacked Ki67 expression in humans (*52*). Interestingly, expression of the transcription factors T-bet and Eomes shares similar kinetics. In mice, T-box transcription factor downregulation and the consequent expression of CD103 on CD8 T cells are essential for TRM formation (*53*). Although the lack of phenotypic changes over time may suggest that airway TRM are a stable effector population, this is not the case since there are major change in gene expression, with genes involved in cytokine production, Th1/Th2 differentiation and the TGFβ pathway highly upregulated at 21 compared to 7 DPI and expression declining at 63 DPI. TGFβ plays a critical role in tissue retention and differentiation into TRM (*54–56*) suggesting that tetramer binding cells at 21 DPI already present TRM features. In addition, we identified different sets of genes characteristic of TRM in humans, which are also upregulated at each of these time points. Others have reported marked differences in gene expression between CD8 T cells specific for different epitopes (*57*), which should be investigated in the future to understand the prevalence of AAV^+^ CD8 T cells.

These data provide a detailed analysis of the phenotype and gene expression of influenza-specific CD8^+^ TRM after influenza infection, but they have not established the protective efficacy of these cells in the pig model. The data imply but do not prove, that the cells are generated in TBLN and migrate via the blood to mucosal sites in the respiratory tract, where we observed a delayed peak response. TRM and circulating memory cells in all sites decline over time, the parallel decline of Ki67 staining suggests that they are not replenished by cell division. Our study was limited to CD8 T cells but future work will need to establish if other memory populations (CD4 and B cells) decline with similar kinetics. Neither have we established, as have others in the mouse, whether TRM in mucosal sites contribute to the recirculating pool of memory cells (*10*). However the extremely high frequency of antigen-specific T cells in BAL and lower frequency in PBMC three weeks after S-FLU immunization, indicate that this contribution may be relatively modest. As aerosol immunization with S-FLU does not induce neutralizing antibodies, it will also be of great interest to determine the persistence of BAL TRM and their role in reducing lung pathology after homologous and heterologous challenge (*23, 25, 58*).

In summary, we have defined the spatial and temporal dynamics and hierarchy of NP-specific responses in a natural host for influenza infection. These data suggest that an airway population of CD8 TRM is established by clonal expansion in draining TBLN and migration to the lungs. Importantly there is no evidence that respiratory tract CD8 TRM decline more rapidly than antigen specific CD8 T cells in other sites. Despite exhibiting a stable phenotype over time, BAL TRM undergo major changes in gene expression. The similarities in phenotype and transcriptional profile of porcine and human TRM highlight the value of this large animal model for understanding the importance of specificity of response in establishing protection to respiratory infections and the development of next generation vaccines.

## Materials and Methods

### Animals and influenza challenge experiment

Animal experiments were conducted according to the UK Government Animal (Scientific Procedures) Act 1986 at the University of Bristol, the Pirbright Institute (TPI) and the Animal and Plant Health Agency (APHA) under project license P47CE0FF2, with approval from ethics committees at each institute. All institutions conform to the ARRIVE guidelines. All pigs were screened prior to experiments for the presence of anti influenza antibody by HAI.

Three influenza challenge experiments were performed at the University of Bristol (T1, T2 and T3), as previously reported (*22*) while a longer time course study (T4) took place at APHA (**Fig. 2 A**). For T1, T2 and T3 thirty-eight Babraham inbred pigs (9.3 weeks old on average) were experimentally infected intranasally, using a mucosal atomisation device (MAD) (MAD300, Wolfe-Tory Medical), with 1 x 10^7^ PFU of MDCK grown H1N1 A/swine/England/1353/2009 (H1N1pdm09), 2ml per nostril. In T1 and T2 one infected pig was culled each day on days 1 to 7, 9, 11 and 13 post challenge while two control uninfected pigs were sampled prior to infection and two more at day 8. During T3 three challenged animals were culled at days 6, 7, 13, 14, 20 and 21 post infection. Six control uninfected animals were included in T3: three culled at day −1 and the other three on the day of challenge. During T4 twelve pigs (10.2 weeks old on average) were challenged as described above and four pigs were culled at each of day 21, 42 and 63 post infection.

A first immunization experiment was performed at TPI where three Babraham pigs of 5.5 weeks of age were sedated and immunized with H7N1 S-FLU [eGFP/N1(PR8)] H7t(Netherlands/219/2003) (2.4. x 10^8^ 50% tissue culture infective dose (TCID50)) administered by aerosol using a SOLO vibrating mesh nebuliser (Aerogen ltd.) as previously described (*23*). A group of two control pigs was left untreated. Vaccinated pigs were boosted after 3 weeks as described above and culled 3 weeks post boost (WPB). Ten minutes prior to sacrifice animals were infused intravenously (i.v.) with anti-CD3 monoclonal antibody (mAb) (clone PPT3), produced in house, at a concentration of 1mg/kg to label circulating T cells. A second immunization experiment took place at APHA. Six Babraham pigs received H1N1 S-FLU [eGFP*/N1(A/Eng/195/2009)] H1(A/Eng/195/2009) (7.3 x 10^7^ TCID50) by aerosol twice, three weeks apart, as described before (*23*). Five unimmunized pigs were used as control. At 3WPB, all animals were challenged intranasally with 2.8 x 10^6^ PFU of H1N1pdm09 delivered by MAD and culled four days later. Half of the pigs in each group received anti-CD3 mAb i.v. 10 minutes prior to culling.

### Tissue sampling and processing

Peripheral blood mononuclear cells (PBMC), tracheo-bronchial lymph nodes (TBLN), lung, broncho-alveolar lavage (BAL) and spleen were processed as previously described (*25, 58*). In addition, nasal turbinate (NT) and trachea were isolated in studies T3 and T4 as described before (*23, 59*). Cells were cryopreserved in 10% DMSO (Thermo Fischer) in fetal bovine serum (FBS).

### IFN**γ** ELISpot assay and identification of influenza nucleoprotein epitopes

Cryopreserved lymphocytes were stimulated using H1N1pmd09 (MOI=1), medium and H1N1pmd09 nucleoprotein derived peptides (GL Biochem Ltd.) and frequencies of IFNγ spot forming cells were determined as described before (*58*). To identify CD8 T cell epitopes in H1N1pdm09 nucleoprotein pools of ten 18 amino acid (aa) peptides overlapping by 12 aa, were used for initial ELISpot screening. Individual peptides from responding pool were then used to identify the highest responding peptides within each pool. Minimal epitopes were then defined using a second screen with 9 aa peptides derived from the 18 aa peptides giving the highest responses (**Supplementary Fig. 2**). Pool 3 and 4 consistently showed the highest responses across tissues and therefore were broken down to identify minimal epitopes of 9 aa length: NP_181-189 AAVKGVGTI_ (AAV) and NP_217-225 VAYERMCNI_ (VAY), which were confirmed to be CD8 epitopes (data not shown) and loaded into porcine SLA-2 molecules to generate tetramers.

### Flow cytometric analysis and cell sorting

Cryopreserved single cell suspensions from PBMC, TBLN, lung, BAL, NT and trachea were thawed and rested for 2 hours in RPMI supplemented with Glutamax, 1% Penicillin-Streptomycin, 5% HEPES and 10% FBS (all from Thermo Fisher) at room temperature before staining. Two million cells per well were stained in 96 well plates with each NP tetramer (AAV, DFE (NP_290–298 DFEREGYSL_) and VAY) separately (as previously described (*23, 32*)). Following tetramer staining, fluorochrome-conjugated antibodies (see **Table S2** for all antibodies used) were added in staining buffer (PBS+0.1% FBS), for 15 mins at 4°C and then washed twice with PBS before fixation with 4% paraformaldehyde solution in PBS (Santa Cruz Biotechnology). Intracellular staining for the detection of Eomesodermin, T-bet and Ki67 was performed using True-Nuclear™ Transcription Factor Buffer Set (BioLegend), according to the manufacturer instructions. Perforin and cytokine staining was achieved using Fixation/Permeabilization Solution Kit (BD Biosciences). Gating strategies for the different panels and control used are illustrated in **Supplementary Fig. 7A and B**. Naïve samples were also stained and % of tetramer^+^CD8 T cells reported in **Table S3**. Stained cells were analyzed using a BD LSRFortessa.

To analyze the time course of activation of different CD8 subpopulations, cryopreserved PBMC from naïve animals were first stained for CD8β, CD45RA and CCR7 expression and sorted into 4 subpopulations (CD45RA^+^CCR7^+^, CD45RA^+^CCR7^-^, CD45RA^-^ CCR7^+^, CD45RA^-^CCR7^-^) using a BD FACSAria cell sorter. Cells were then centrifuged at 1000 x g for 5 minutes and re-suspended in cell culture medium (RPMI, 10% FBS, HEPES, Sodium pyruvate, Glutamax and Penicillin/Streptomycin) overnight at 37°C. On the following day, cells were stimulated using PMA Ionomycin (BioLegend) for 2, 4 and 6 hours and cytokines detected by intracellular cytokine staining (see **Table S2** for antibodies list) using Fixation/Permeabilization Solution Kit (BD Biosciences), unstimulated cells were used as a control.

Cells isolated from BAL, TBLN and PBMC were stimulated with H1N1pdm09 (MOI=1) overnight and cytokine production quantified by intracellular cytokine staining as previously described (*22*). Media only controls for each sample were used as baseline and subtracted from the results.

For preparation of RNA, BAL cells isolated from 7, 21 and 63 DPI samples (n=3) were depleted of alveolar macrophages using a MACS cell separation LS column (Miltenyi Biotec) after staining for CD14 and CD172a (see **Table S2**). Lymphocytes were then stained for DFE tetramer, CD8 and live/dead marker using a BD FACSAria (BD Bioscience).

Data were analyzed using FlowJo software v10.7 (Tree Star).

### RNA-seq

BAL DFE^+^ T cells were sorted (average of 4480 cells/sample) in PBS, samples were then centrifuged at 3000g for 5 minutes. RNA was extracted using PicoPure RNA Isolation Kit (Thermo Fisher) according to the manufacturer instructions followed by DNase treatment (TURBO DNA-free™ Kit, Thermo Fisher). Isolated RNA was used as input for SMARTer Stranded Total RNA-Seq Kit v3 - Pico Input Mammalian (Takara Bio) and PCR amplification performed. cDNA was pooled and sequenced on NovaSeq using an S1 100 PE flow cell. Raw fastq files were used for an initial quality control using FastQC (https://www.bioinformatics.babraham.ac.uk/projects/fastqc/). Next, CogentAP (Cogent NGS Analysis Pipeline v1.0, Takara Bio) was used to trim and add the sample barcodes to the fastq header for each sample. The reads were then trimmed of Illumina and library prep adapters using cutadapt (*60*), and subsequentially aligned to Sus scrofa Sscrofa11.1 assembly (GCA_000003025.6) using STAR (*61*) with default parameters. UMI-tools was used to discard duplicated reads (*62*) and featureCounts extracted the number of reads aligned to each gene feature (*63*). Differential gene expression (DGE) analysis was carried out using TCC-GUI, which iterates DESeq2 for data normalisation, using the featureCounts output for pairwise comparisons (*64, 65*). Pathway analysis was performed using Webgestalt online tool (*66*), selecting GSEA as comparison method and sscrofa as reference genome.

KEGG pathway database was used as reference and ensemble ID were uploaded together with ranked score (-log_10_(p value)*log_2_(fold change difference)) based on the results of DGE comparison. Other settings include: minimum number of IDs in the category (5), maximum number of IDs in the category (2000), significance level (FDR < 0.05) and number of permutation (1000).

## Statistical analysis

To compare the changes over time in the proportion of CD8^+^ T cells specific for each tetramer in a tissue, the following curve was fitted to the data, namely,

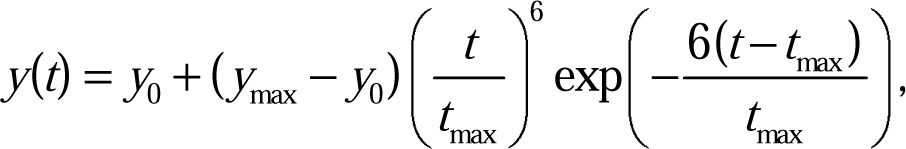

where y is the proportion at t days post infection, y0 is the baseline proportion, ymax is the peak proportion and tmax is the time of peak proportion. Estimated parameters for each tetramer and tissue are listed in **Table 1**. This curve was chosen as it gives an appropriate shape for the data without including a large number of parameters.

Similarly, changes over time in the proportion of tetramer-specific T cells expressing different cell surface markers in each tissue were compared by fitting exponential curves to the data,

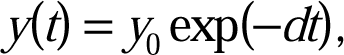

where y is the proportion at t days post infection, y0 is the initial proportion and d is the decay rate (/day). Each marker was analyzed independently.

In both analysis, variation in parameters (i.e. y_0_, y_max_ and t_max_ or y_0_ and d) amongst tissues and tetramers was assessed by fitting different models to the data by nonlinear least squares and comparing the residual deviance for the models using F-tests (*67*). These analysis were implemented in Matlab (version R2020b; The Mathworks, Inc.).

Because data on CD8^+^ T cells in the trachea were only available for a reduced number of time points, the proportion of cells specific for each tetramer in this tissue were compared at each time point using a Kruskal-Wallis test followed by pairwise Wilcoxon rank-sum tests. This analysis was implemented in R (version 4.0.5) (*68*).

To assess changes over time in the relative proportion of CD8^+^ T cells specific for each tetramer, the following model was used:

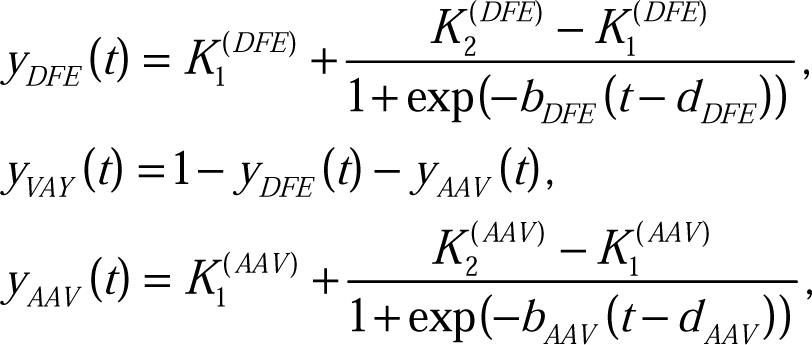

where y(t) is the proportion specific for the tetramer t days post infection, K1 and K2 are the minimum and maximum frequencies, b the rate of change in frequency and d is the time of the maximum rate of change. This formulation ensures the total proportion of cells is 100%. Variation in parameters (i.e. K_1_, K_2_, b and d) amongst tissues was assessed by fitting different models to the data by nonlinear least squares and comparing the residual deviance for the models using F-tests (Ross 1990) (**Table S4**). This analysis was implemented in Matlab (version R2020b; The Mathworks, Inc.). Only samples for which the frequency of all three tetramers was available were included in this analysis.

Trends in the phenotype of tetramer specific CD8^+^ T cells in each tissue were assessed using linear models. Each model included the proportion of cells in the population (CD45RA^+^/^-^ and CCR7^+^/^-^) specific for a tetramer as the response variable and tissue, tetramer and days post infection as explanatory variables, together with two- and three-way interactions between the explanatory variables. Model simplification proceeded by stepwise deletion of non-significant (P>0.05) terms (as judged by F-tests). This analysis was implemented in R (version 4.0.5) (R Core Team 2021).

Frequencies of cytokine secreting cells in DFE^+^ and AAV^+^ cells over time were compared using two-way ANOVA in GraphPad Prism version 9.1.0.

## Supplementary Materials

### Supplementary Figures

**Supplementary Figure 1. Expression of perforin and CD27 in CD8 T cell subset**

**Supplementary Figure 2. Identification of NP epitopes AAV and VAY**

**Supplementary Figure 3. Number of tetramer^+^ T cells in tissues**

**Supplementary Figure 4. Distribution of tetramer^+^ cells in the trachea**

**Supplementary Figure 5. Gene expression of tissue resident memory T cells features.**

**Supplementary Figure 6. CD3 infusion for the identification of tissue resident memory T cells.**

**Supplementary Figure 7. Gating strategy and controls.**

### Supplementary Tables

**Table S1.** Relevant significant (FDR<0.05) KEGG pathways upregulated in 21DPI vs 7DPI comparison.

**Table S2.** List of antibodies used.

## Acknowledgments

We are grateful to all animal staff at the University of Bristol, the Pirbright Institute and APHA. We thank Emily Porter and the Mucosal Immunology group at TPI for their help in tissue collection and processing. We acknowledge the Bioimaging group at TPI for their support with cell sorting and Edinburgh Genomics for their help with RNA sequencing.

## Author contributions

Conceptualization: ET, VM, AT, PB

Methodology: VM, ME, SG, SJ, BP, AM, SM, TM, PR, RI, AKS

Investigation: VM, ET, SG, SJ, TC Visualization: VM, SG

Funding acquisition: ET, BC, AT Supervision: ET, AT, PB

Writing – original draft: VM, ET, PB

Writing – review & editing: VM, ET, PB, ME, RI, TC, AKS, BC

## Data and material availability

RNA-seq data will be available on GEO website after manuscript submission.

