## Supplementary material for "Spatial, temporal and molecular dynamics of swine influenza virus-specific CD8 tissue resident memory T cells": SupFig1.pdf

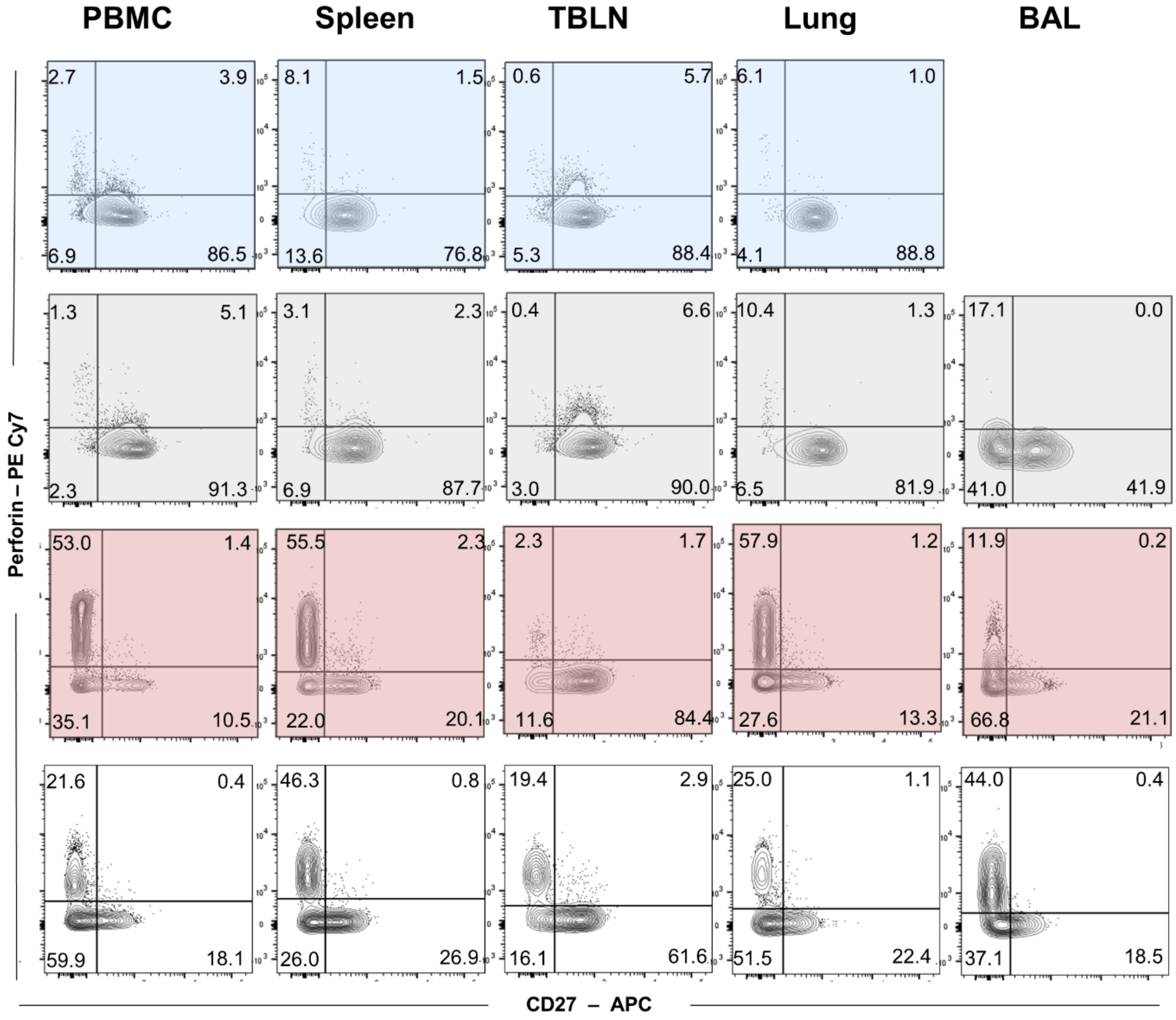


**Supplementary Figure 1. Expression of perforin and CD27 in CD8 T cell subset**

Representative plots showing perforin and CD27 expression in CD8 T lymphocyte subsets defined by CD45RA and CCR7 expression: naïve (CD45RA$^+$, CCR7$^+$, blue), TCM (CD45RA$^-$, CCR7$^+$, grey), TEM (CD45RA$^-$, CCR7$^-$ , red panels) and TDE (CD45RA$^+$ CCR7$^-$, white panels). Each quadrant shows the mean percentage of 3 animals.
