## Supplementary material for "Spatial, temporal and molecular dynamics of swine influenza virus-specific CD8 tissue resident memory T cells": SupFig2.pdf

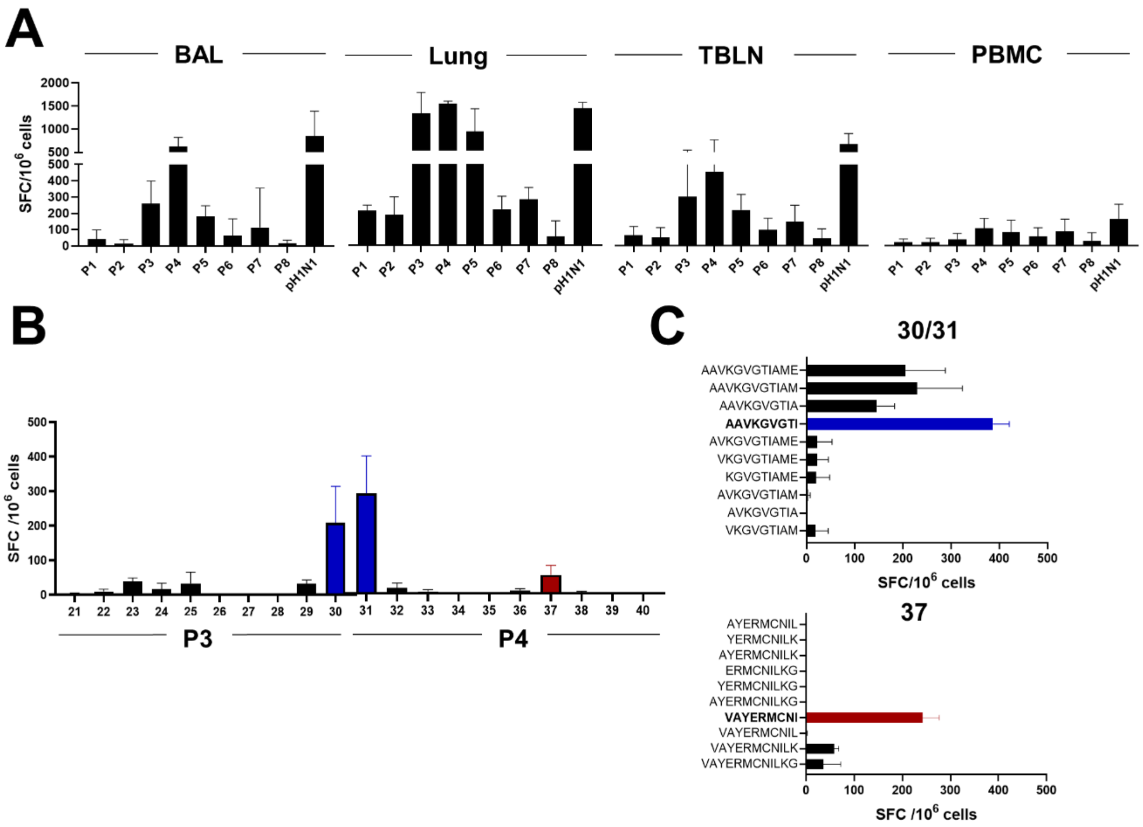


**Supplementary Figure 2. Identification of NP epitopes AAV and VAY (A)** Pools of 10 peptides of 18 amino acid (aa) length from the NP of H1N1pdm09 were used for an initial screen to stimulate cells isolated from BAL, lung, TBLN and PBMC at 13/14 DPI. Responses were measured by IFNg ELISpot and H1N1pdm09 (pH1N1) MOI = 1 was used as a positive control. **(B)** Spot forming cells (SFC) in TBLN after stimulation with individual peptides from pool 3 (p3) and pool 4 (p4), highlighted the peptides that subsequentially identified AAV epitope (in blue) and VAY (in red). **(C)** Minimal epitope identification  (using peptides of 9 aa) from peptide 30/31 (top) and 37 (bottom) identify the AAV (in blue) and VAY (in red) epitopes. Mean and SEM of 3 animals.
