## Supplementary material for "Spatial, temporal and molecular dynamics of swine influenza virus-specific CD8 tissue resident memory T cells": SupFig3.pdf

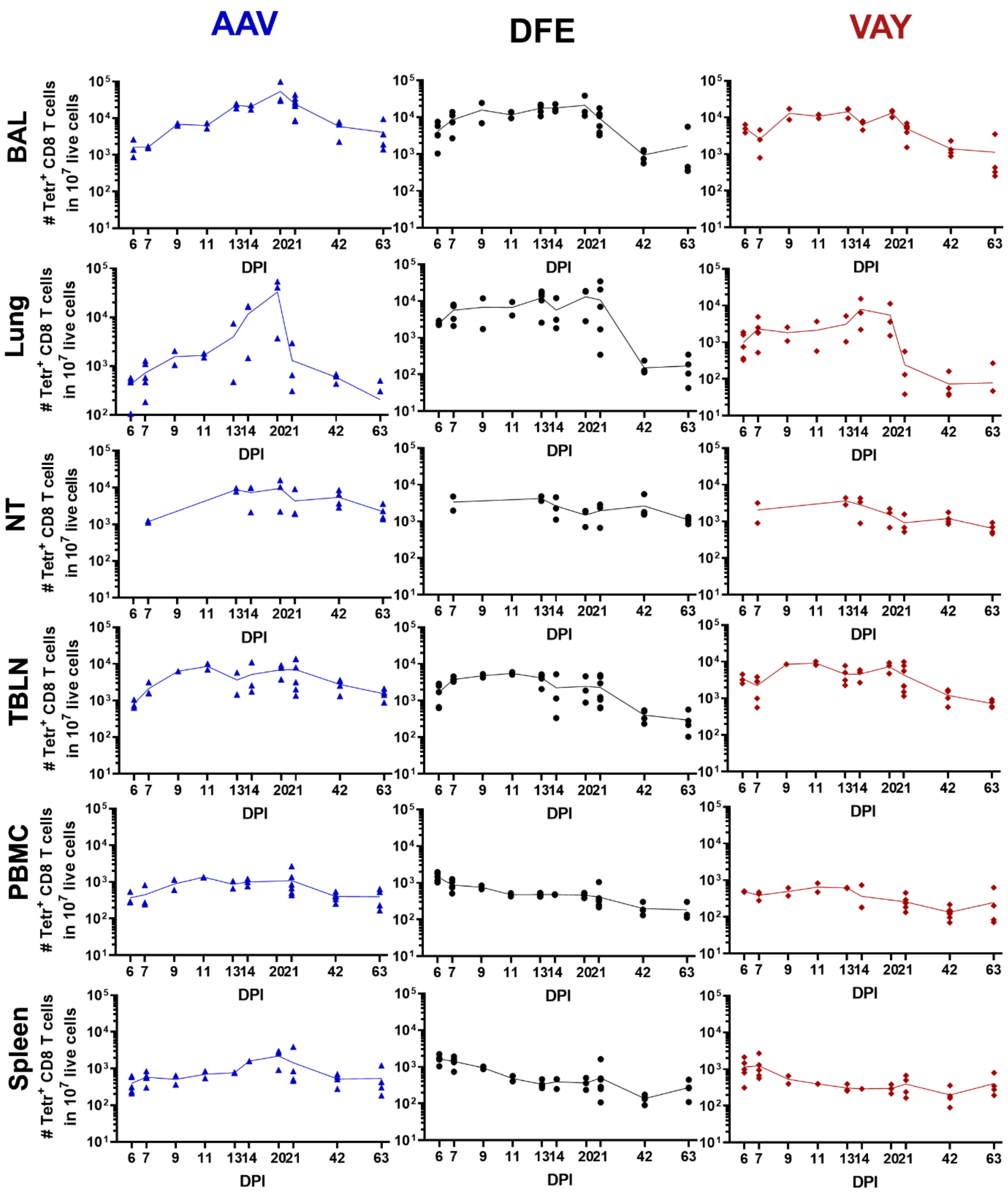


**Supplementary Figure 3. Number of tetramer⁺ T cells in tissues** Counts of tetramers⁺ CD8 T cells in 10 million live cells isolated from each tissues. The symbols indicates individual animals while lines connect the mean at each timepoint.
