## Supplementary material for "Spatial, temporal and molecular dynamics of swine influenza virus-specific CD8 tissue resident memory T cells": SupFig4.pdf

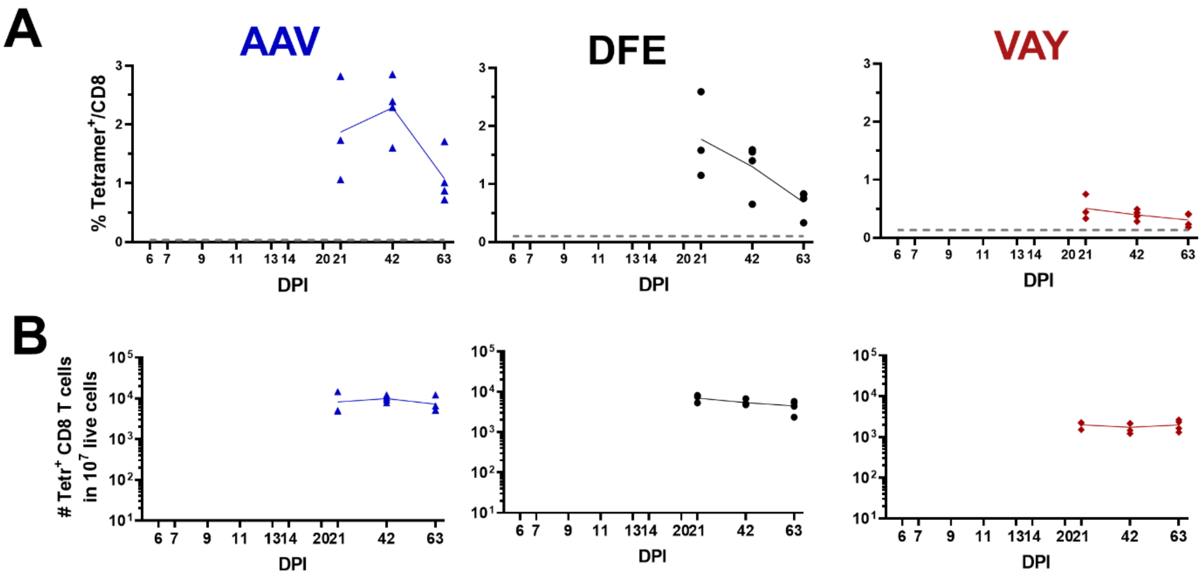


**Supplementary Figure 4. Distribution of tetramer+ cells in the trachea (A)** Percentages of DFE, VAY and AAV+ within CD8 T cell population at day 21, 42 and 63 post infection **(B)** Number of tetramer+ T cells in 10 million live cells. Each symbol represent an individual and the dotted line the average % of tetramer+ T cells in naïve animals.
