## Supplementary material for "Spatial, temporal and molecular dynamics of swine influenza virus-specific CD8 tissue resident memory T cells": SupFig5.pdf

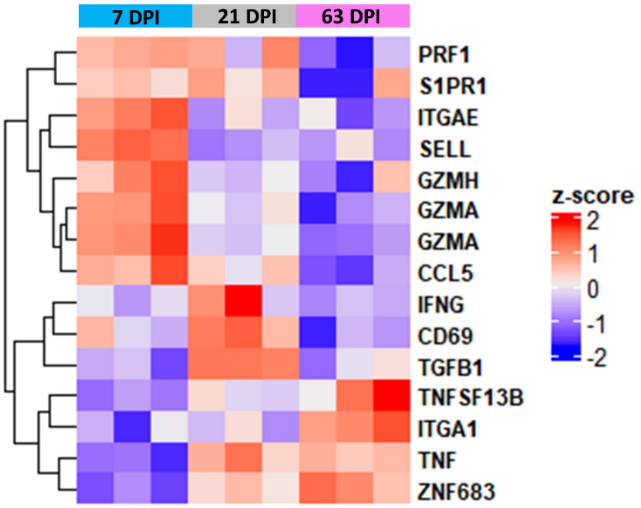


**Supplementary Figure 5. Gene expression of tissue resident memory T cells features.** Heatmap of selected genes related to tissue resident memory T cells.
