## Supplementary material for "Spatial, temporal and molecular dynamics of swine influenza virus-specific CD8 tissue resident memory T cells": SupFig6.pdf

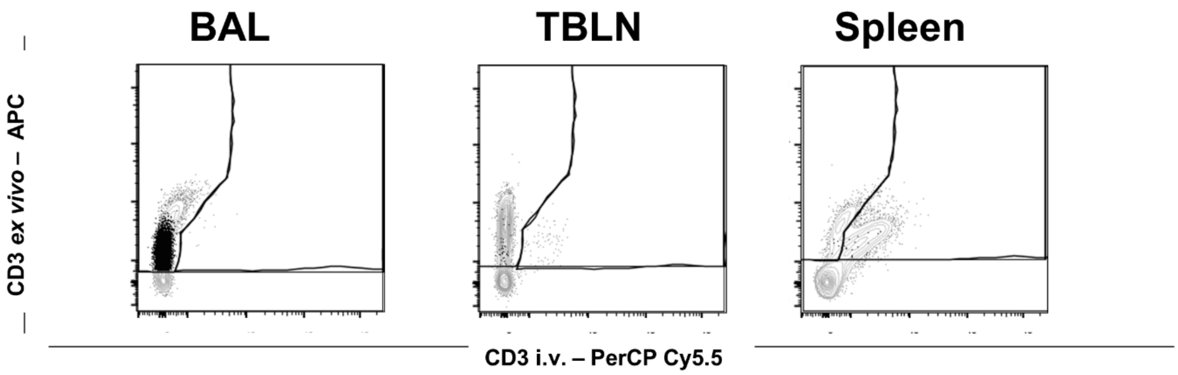


**Supplementary Figure 6. CD3 infusion for the identification of tissue resident memory T cells.** Representative FACS plots showing CD3 staining of DFE$^{+}$ T cells (in back) and total live lymphocytes (in grey) in BAL, TBLN and spleen of S-FLU aer immunized animal 3 weeks post boost.
