## Supplementary material for "Spatial, temporal and molecular dynamics of swine influenza virus-specific CD8 tissue resident memory T cells": SupFig7.pdf

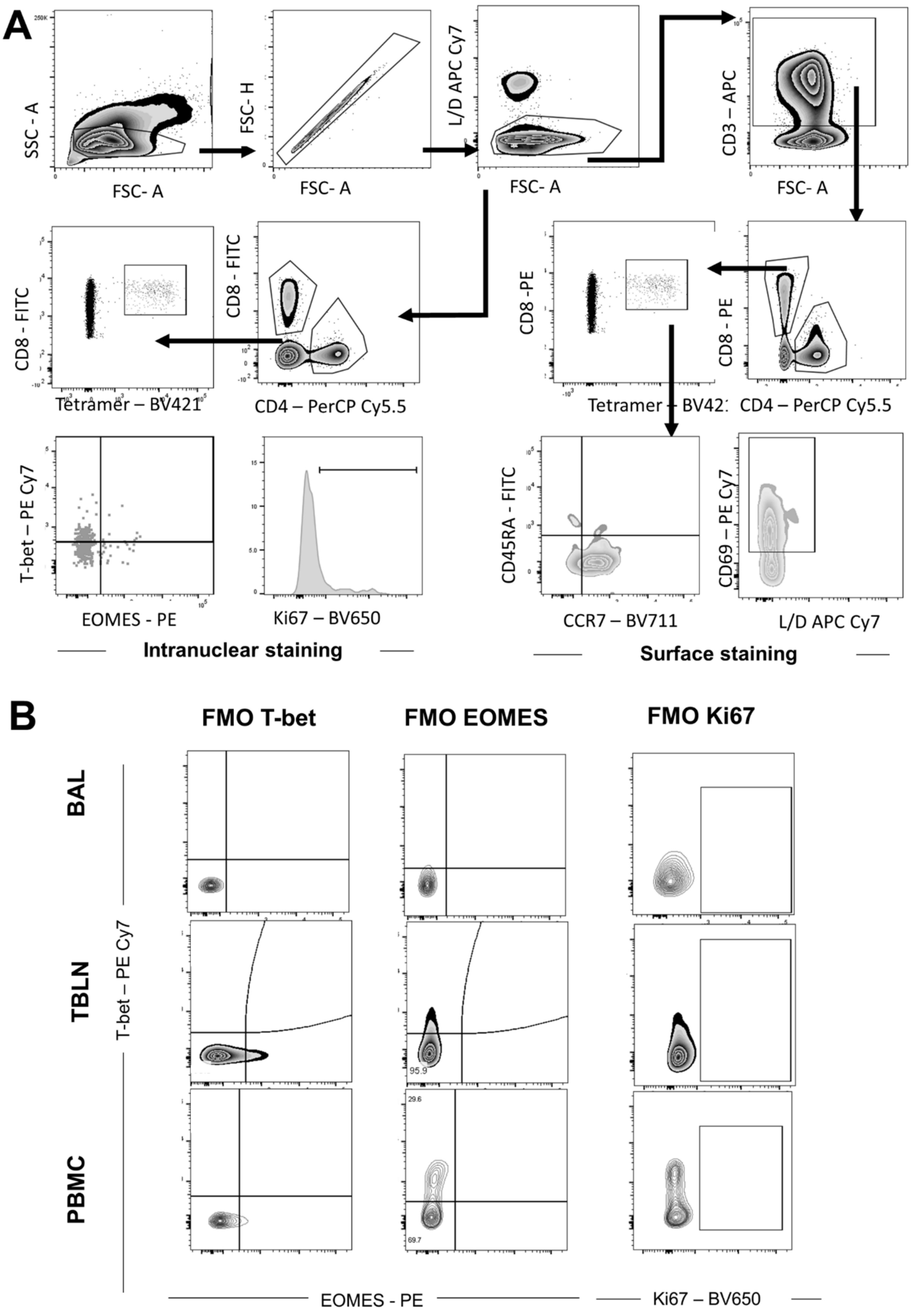

A
SSC - A
FSC- A
FSC- H
FSC- A
L/D APC Cy7
FSC- A
CD3 – APC
FSC- A
CD8 - FITC
Tetramer – BV421
CD8 - FITC
CD4 – PerCP Cy5.5
CD8 -PE
Tetramer – BV421
CD8 - PE
CD4 – PerCP Cy5.5
T-bet – PE Cy7
EOMES - PE
Ki67 – BV650
CD45RA - FITC
CCR7 – BV711
CD69 – PE Cy7
L/D APC Cy7
Intranuclear staining
Surface staining
B
FMO T-bet
FMO EOMES
FMO Ki67
BAL
TBLN
PBMC
T-bet – PE Cy7
95.9
29.6
69.7
EOMES - PE
Ki67 – BV650

**Supplementary Figure 7. Gating strategy and controls. (A)** Representative FACS plots (from PBMC) showing the gating strategy for intranuclear staining (panels on the left) and surface marker staining (on the right.) **(B)** Fluorescence minus one (FMO) controls for transcriptional factors in the different tissues
