## Supplementary material for "Spatial, temporal and molecular dynamics of swine influenza virus-specific CD8 tissue resident memory T cells": TableS1.pdf

**Table S1.** Relevant significant (FDR<0.05) KEGG pathways upregulated in 21DPI vs 7DPI comparison.

| Gene Set | Pathway | Normalised enrichment score | P Value | FDR |
|---|---|---|---|---|
| **ssc04640** | Hematopoietic cell lineage | 0.90817 | 0 | 0 |
| **ssc04010** | MAPK signalling pathway | 0.77936 | 0 | 0 |
| **ssc04658** | Th1 and Th2 cell differentiation | 0.88885 | 0 | 0 |
| **ssc05164** | Influenza A | 0.82988 | 0 | 0 |
| **ssc04668** | TNF signalling pathway | 0.85347 | 0.0014388 | 0.0014388 |
| **ssc04060** | Cytokine-cytokine receptor interaction | 0.75969 | 0 | 0 |
| **ssc04612** | Antigen processing and presentation | 0.88222 | 0 | 0 |
| **ssc04350** | TGF-beta signalling pathway | 0.83065 | 0.0014925 | 0.0014925 |
