## Supplementary material for "Spatial, temporal and molecular dynamics of swine influenza virus-specific CD8 tissue resident memory T cells": TableS2.pdf

**Table S2.** List of antibodies used.

| Antigen | Clone | Isotype | Fluorochrome | Source of primary Ab | Details of secondary Ab |
|---|---|---|---|---|---|
| CD4 | 74-12-4 | IgG2b | PerCP-Cy5.5 | BD Biosciences | |
| CD8β | PPT23 | IgG1 | FITC | Bio-Rad Laboratories | |
| CD8β | PPT23 | IgG1 | PE | Bio-Rad Laboratories | |
| TNF | MAb11 | IgG1 | BV421 | BioLegend | |
| IFNγ | P2G10 | IgG1 | APC | BD Biosciences | |
| IL-2 | A150D 3F1 2H2 | IgG2a | PE-Cy7 | ThermoFisher | rat-anti-mouse, IgG2a, BioLegend |
| CCR7 | 3D12 | IgG2a | BV711 | BD Biosciences | |
| CD45RA | MIL13 | IgG1 | FITC | Bio-Rad Laboratories | |
| Annexin V | N/A | N/A | BV510 | BioLegend | |
| CD69 | 01-14-22-51 | IgG2b | PE Cy7 | Kyoto Institute of Nutrition & Pathology (Hayashi et al. 2018) | Goat-anti-mouse, BioLegend |
| EOMES | WD1928 | IgG1 | PE | eBioscience | |
| T-bet | eBio4B10 | IgG1 | PE Cy7 | eBioscience | |
| Ki67 | B56 | IgG1 | BV650 | BD Biosciences | |
| Perforin | δG9 | IgG2b | Purified | | Lightning-Linked PE-Cy7, antibody labelling kit, Novus Bio |
| CD27 | B30C7 | IgG1 | APC | Bio-Rad Laboratories | |
| CD14 | REA599 | IgG1 | PE | Miltenyi Biotec | |
| CD172a | BL1H7 | IgG1 | PE | Bio-Rad Laboratories | |
| Live/Dead Marker | N/A | N/A | Fixable Near-IR | Life Technologies | |
